# Targeted memory reactivation during sleep elicits signals related to learning content

**DOI:** 10.1101/449389

**Authors:** Boyu Wang, James W. Antony, Sarah Lurie, Paula P. Brooks, Ken A. Paller, Kenneth A. Norman

## Abstract

Reactivation of learning-related neural activity patterns is thought to drive memory stabilization. However, finding reliable, non-invasive, content-specific indicators of reactivation remains a central challenge. Here, we attempted to decode the content of reactivated memories in the electroencephalogram (EEG) during sleep. During encoding, human participants learned to associate spatial locations of visual objects with left- or right-hand movements, and each object was accompanied by an inherently related sound. During subsequent slow-wave sleep within an afternoon nap, we presented half of the sound cues that were associated (during wake) with left- and right-hand movements before bringing participants back for a final post-nap test. We trained a classifier on sleep EEG data (focusing on lateralized EEG features that discriminated left- vs. right-sided trials during wake) to predict learning content when we reactivated the memories during sleep. Discrimination performance was significantly above chance and predicted subsequent memory, supporting the idea that reactivation leads to memory stabilization. Moreover, these lateralized signals increased with post-cue spindle power, demonstrating that reactivation has a strong relationship with spindles. These results show that lateralized activity related to individual memories can be decoded from sleep EEG, providing an effective indicator of offline reactivation.

## Introduction

In recent decades, evidence has increasingly converged on the idea that newly formed memory traces are spontaneously reactivated during sleep, promoting their stabilization (Wilson and McNaughton, 1994; Ji and Wilson, 2007). Decoding the content of replayed information from animal studies (using invasive electrophysiology) has massively advanced our understanding of how sleep contributes to learning and memory. Detecting non-invasive, content-specific signals of reactivation in humans is, therefore, of utmost interest for understanding brain mechanisms of memory.

Several recent studies have brought us closer to this goal. One study used multivariate classification methods to decode (during sleep) whether people had viewed large numbers of faces or scenes prior to sleep (Schönauer et al., 2017). However, as this study queried spontaneous reactivation of category (face vs. scene) information, it was not possible to determine which individual face and scene memories were being reactivated. One way to gain some control over which memories are reactivated is to play auditory cues associated with specific studied stimuli during sleep; one can then relate any evoked neural patterns to memory for the specific stimuli that were cued. This approach, termed targeted memory reactivation (TMR; Oudiette & Paller, 2013), has been shown to bias neuronal replay in the hippocampus in rodents (Bendor and Wilson, 2012) and benefit memory in humans (Rasch et al., 2007; Rudoy et al., 2009), suggesting it resembles reactivation that occurs spontaneously.

Recently, Cairney et al. (2018) used TMR to demonstrate reactivation of specific episodic associations. Their participants learned pairings of auditory words with either face or scene images; effectively, the category of the associated image served as an item-specific episodic “tag” for each word. Cairney et al. used the words as TMR cues during sleep, and found that the category of the associated image (face or scene) could be successfully decoded during post-TMR cue activity (Cairney et al., 2018). Additionally, Cairney et al. explored the relationship between content decoding and sleep spindles, which are brief (0.5 - 3 s) bursts of 11-16 Hz oscillatory brain activity during NREM sleep that frequently correlate with memory retention (Antony and Paller, 2017). Cairney et al. found that the time intervals when face/scene content decoding was possible overlapped with the window when sleep spindles were present.

Here, we explored whether we could leverage lateralized neural signals as a “tag” to track reactivation of specific episodic associations during sleep. Several prior findings suggest that lateralized neural signals can be tracked with EEG. After training participants to perform lateralized judgments on words during wake (e.g., “press left if word, press right if nonword”), corresponding lateralized responses were elicited by those stimuli during sleep (Kouider et al., 2014; Andrillon et al., 2016). This finding shows that participants can apply a well-learned task set (acquired during wake) to stimuli presented during sleep, but does not show retrieval of specific episodic memories. Additionally, lateralized differences in the number and amplitude of fast sleep spindles emerged after reactivating learning contexts using learning-related odors that were predominantly associated with left- or right-sided visual stimuli (Cox et al., 2014). This finding, like the Schonauer et al. (2017) finding noted above, shows that participants can reactivate “tags” associated with a learning context (here, left vs. right stimuli), but does not indicate which memories within that context are being reactivated.

In the present study, we sought to extend the aforementioned results by “tagging” individual declarative memories with lateralized movements, reactivating them during sleep via auditory TMR cues, and looking for reactivation evidence via lateralized neural activity. Participants (*N*=24) first encoded visual objects in one of four locations on the left or right side of a computer touch screen (Fig 1). Half of the objects were designated targets. Each target (e.g. cow) was accompanied by an inherently related sound (e.g. [moo]) and was presented concurrently with a visual non-target (e.g. avocado) on the opposite side of the screen. Each non-target was unrelated to the sound. During encoding, participants were instructed to imagine moving their corresponding hand towards the target (e.g., right hand for right-side stimuli) for five seconds, after which they executed the movement. We predicted that participants would show significant lateralized activity for left- and right-side target trials during the imagination period. Participants then took an item-location test for each target without feedback before an afternoon nap. During indications of slow-wave sleep (SWS), we presented half of the sounds (i.e. TMR cues) that were previously associated with both left- and right-side targets. We reasoned that neural features that discriminated between these trial types while awake could be used to decode neural activity evoked by TMR cues during sleep. After a 90-min break, participants returned for a final memory test.

**Figure 1.**
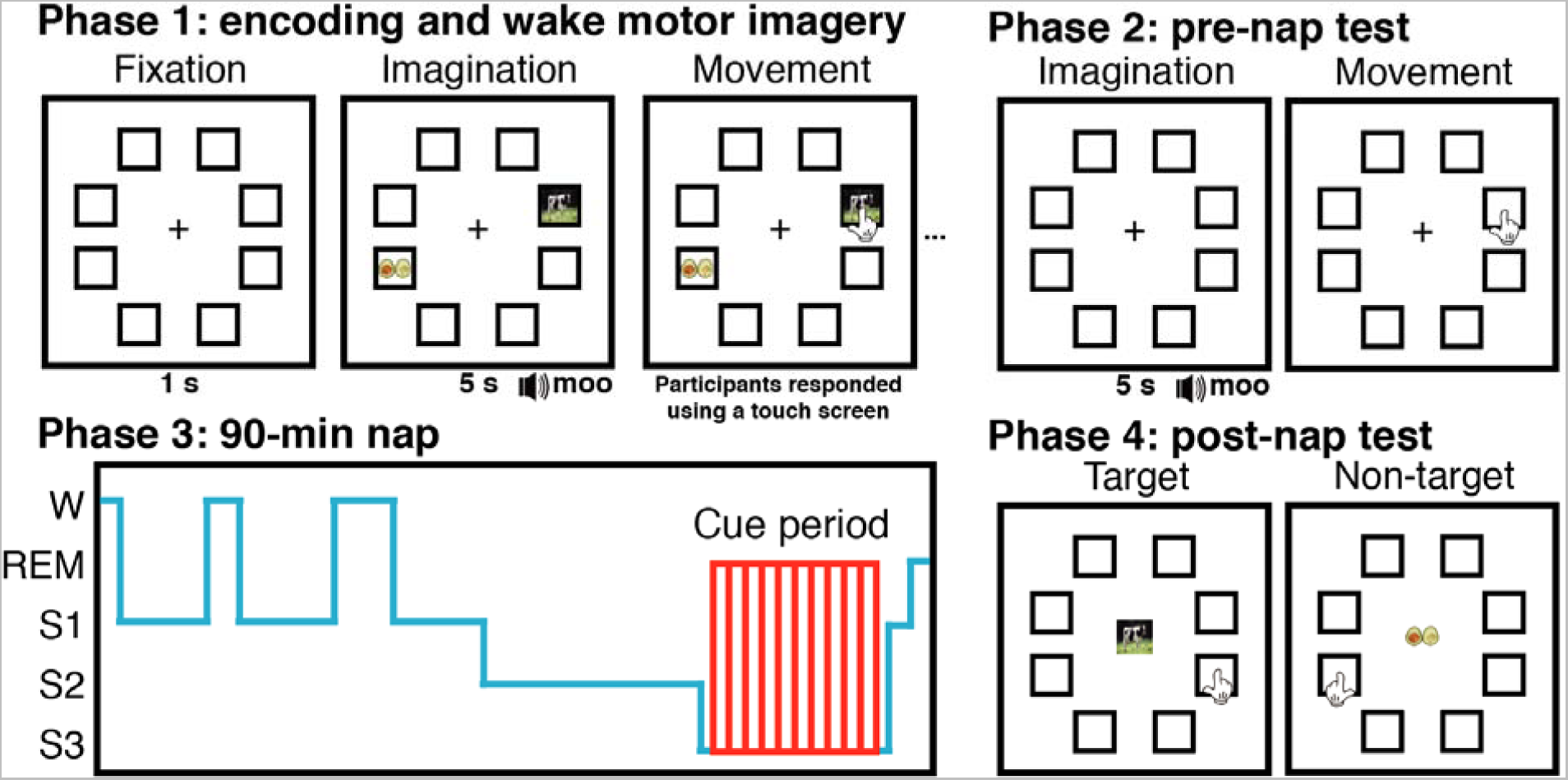
Experimental task and design. Participants imagined reaching out and touching targets that were associated with auditory cues before executing their movement. Participants used either their left or right hand depending on the location of the target on the screen (left hand for left-side targets, right hand for right-side targets). Participants then took a pre-nap test on the location of each target. During periods of SWS in an afternoon nap, participants were presented with half of the sounds associated with left-side and half associated with right-side targets. After the nap, participants returned to the lab and took a test on all target and non-target locations.

Behaviorally, we predicted participants would remember cued targets better than uncued ones. Neurally, we aimed to detect lateralized, learning-related EEG activity during sleep following TMR cues associated with left- vs. right-side targets. We predicted that these lateralized signals would be larger for targets with intact pre-nap memory and would also predict post-nap memory. We also assessed the relationship between lateralized reactivation evidence and other neural predictors of post-nap memory in the sleep EEG. TMR benefits have been intimately linked with post-cue spindles (Schreiner et al., 2015; Antony et al., 2018a, 2018b). Therefore, we anticipated that post-cue spindles would positively predict lateralized reactivation signals during sleep, as well as subsequent memory during the post-nap test. Furthermore, prior work has shown that spindles have a refractory period of 3-6 s, during which TMR cues are less effective (Antony et al., 2018b); as such, we expected that pre-cue spindles would negatively predict both lateralized reactivation signals and subsequent memory.

## Results

### Behavior

Prior to sleep, participants took an item-location memory test on all targets, with 8 possible locations (chance proportion correct = 0.125). Performance did not differ on the pre-nap test between targets that were later cued (mean proportion correct and standard error of measurement: 0.55 ± 0.024) and uncued (0.55 ± 0.027; *t*(23) = 0.06, *d_z_* = 0.01, *p* = 0.95). After the nap, participants returned to the lab to take another item-location memory test on all targets and non-targets. We measured memory retention for targets by subtracting pre-nap performance from post-nap performance while regressing out pre-nap performance, similar to previous reports (Antony et al., 2018a, 2018b). We measured retention for non-targets by assessing post-nap accuracy, since these were not tested prior to the nap. We predicted that cued targets would show better retention than uncued targets. Contrary to our prediction, this difference was not significant (cued: 0.32 ± 0.02; uncued: 0.29 ± 0.02; *t*(23) = 1.4, *d_z_* = 0.29, *p* = 0.17). However, we reasoned that reactivation may only be possible for items remembered pre-nap since feedback was not given after the pre-nap test. Therefore, we ran another analysis looking at the proportion of items still remembered post-nap that were remembered pre-nap. We found that cued targets were remembered marginally better than uncued targets (cued: 0.85 ± 0.02; uncued: 0.81 ± 0.03; *t*(23) = 1.8, *d_z_* = 0.38, *p* = 0.07). Finally, there was no difference between the proportion of remembered non-targets whose sound was associated with cued targets versus uncued targets (cued: 0.28 ± 0.02; uncued: 0.29 ± 0.02; *t*(23) = 0.63, *d_z_* = 0.13, *p* = 0.53).

### Feature selection via decoding during wake

Our primary goal in analyzing the wake EEG data was to find neural features that discriminated between left and right conditions during motor imagery at encoding, with the aim of using those features for classification during sleep. To accomplish this, we relied on the prominent signal produced by the lateralized readiness potential as a result of imagining motor activity from opposite limbs (Kouider et al., 2014). First, we subtracted each lateralized electrode on the right side of the head from its left side counterpart (e.g., F3 - F4) and eliminated ten midline electrodes (e.g., Fz), reducing the number of independent features from 64 channels to 27 channel pairs. Any activity differing from zero was thus linked with lateralized activity.

Next, as EEG data can be highly correlated across channels, we reduced the dimensionality of our data by running spatial independent components analysis (ICA; Delorme and Makeig, 2004). We combined waking data from all participants and ran spatial ICA to obtain across-subject independent components (Fig 2A). We then asked which components significantly discriminated between the conditions (left vs. right) during wake (see Methods for details). Plots of significant components are shown locked to the visual stimulus onset when participants were directed to begin motor imagery (Fig 2B). These components thus provided neural features of learning-related activity by which we could assess reactivation during sleep.

**Figure 2.**
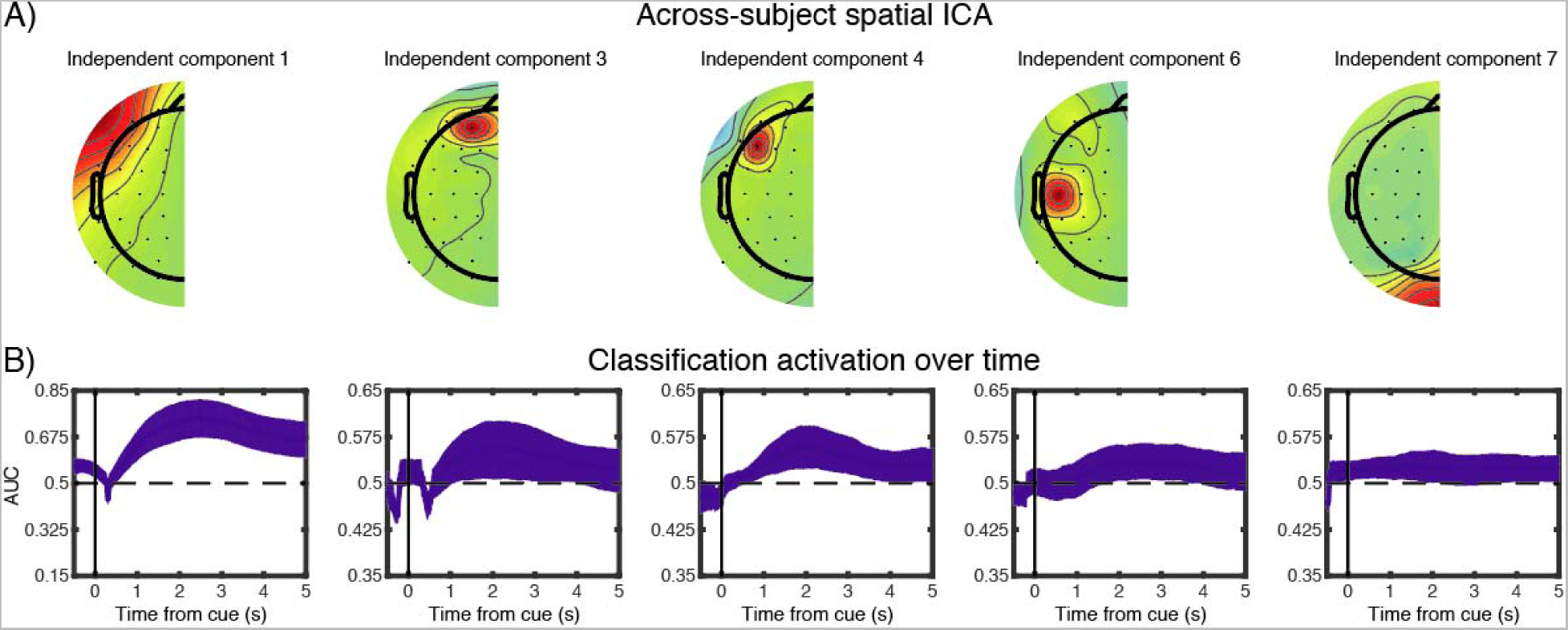
Wake classification and feature selection for sleep. Voltages from lateralized EEG electrodes were subtracted and midline electrodes were eliminated to create 27 electrode pairs. Next, data from all participants were combined; ICA was run on the combined data, and we assessed which of the resulting components discriminated between left and right motor imagery during wake. (A) Shows the five components (1, 3, 4, 6, and 7) that significantly discriminated between left and right motor imagery. (B) Shows classification accuracy (operationalized as AUC, area under the curve) as a function of time bin for those five components. Error bars indicate the edges of the middle 95^th^ percentile of classification accuracies. X-axis values indicate the center time point for each 1000-ms time bin. Note that there is a wider y-axis range for component 1 than other components.

### Content-related activity during sleep differs by pre- and post-nap target memory

We next sought to obtain evidence of learning-content-related activity during sleep. We investigated neural activity after playing TMR cues that were paired with targets associated with left- versus right-side movements (hereafter, left-targets and right-targets). We trained a multivariate classifier on sleep EEG data, using (as input features) the same ICA components that were significantly discriminable during wake (see Methods for details).

We conducted our sleep-classification analyses in three steps. First, we asked whether left-targets and right-targets were discriminable when we allow all items to enter in the analysis. We found the classifier significantly predicted lateralized category, most prominently between 1376 and 4270 ms post-cue (*p* = 0.01; Fig 3A).

**Figure 3.**
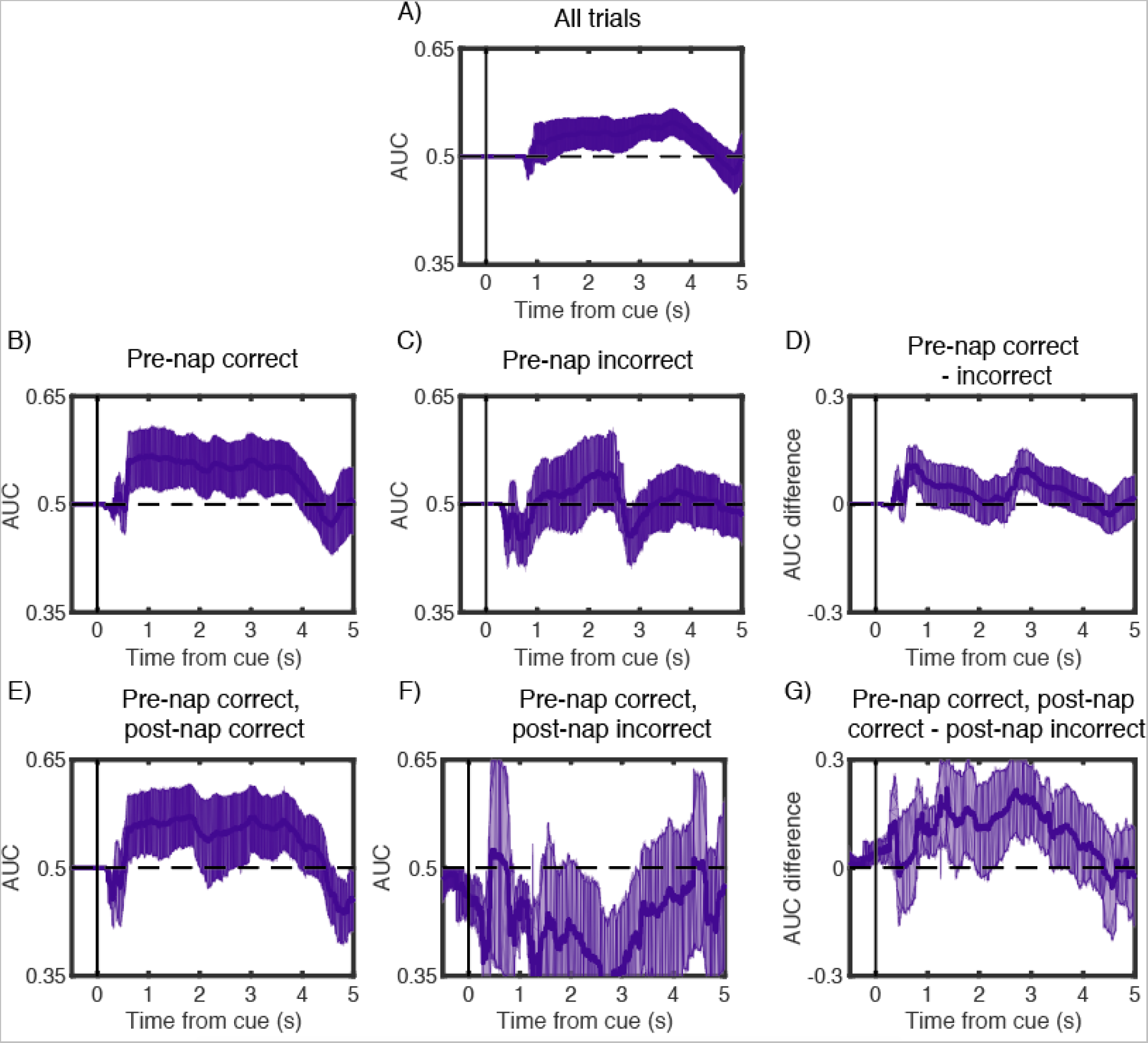
Lateralized, content-related reactivation during sleep related to pre-nap and post-nap memory. Using the lateralized components that discriminated left vs. right motor imagery during wake as features, we trained a leave-one-subject-out classifier on sleep data and computed AUC-based classification performance time-locked to TMR cues. We found significant classification performance when using all trials (A). We also found significant classification performance when focusing just on trials that were remembered pre-nap (B), but not for trials that were not remembered pre-nap (C). However, there were not significant differences in classification performance between these conditions (D). Using only trials that were remembered before the nap, we then found significant classification performance for trials that were subsequently remembered post-nap (E), but not for those forgotten post-nap (F), and there were significant differences in classification performance between these conditions (G). In all graphs, classification was performed on time-averaged signals within 1000-ms windows centered on each timepoint, and error bars indicate the edges of the middle 95^th^ percentile of classification accuracies.

Second, we reasoned that lateralized activity should be greater for targets whose position was correctly remembered before the nap. As there was no feedback given after the pre-nap test, these were the only cues that we could verify were correctly encoded. For items recalled pre-nap, we again found significant classification, in this case most prominently between 592 and 3924 ms post-cue (*p* = 0.01; Fig 3B). Conversely, for items not remembered pre-nap, classification was not significantly predictive (no significant time points, *p* > 0.99; Fig 3C). We also directly contrasted decoding ability for items initially remembered versus those not initially remembered by subtracting the classification accuracy for each condition on each resampled subset, and then assessed whether this differed from what is expected by chance by repeating this analysis using scrambled labels. This difference was not significant (*p* = 0.195; Fig 3D).

Third, limiting ourselves to items that were correctly remembered pre-nap, we further asked whether there were differences in classification based on post-nap memory. For items that were remembered on both the pre- and post-nap tests, we found significant above-chance classification, most prominently from 592 to 2016 ms and 2804 to 3932 ms (*p* = 0.015; Fig 3E). For items remembered only on the pre-nap test but not the post-nap test, we found no meaningful above-chance segments (*p* = 0.605; Fig 3F). Finally, we directly contrasted these signals and found that classification was significantly higher for items remembered pre-nap and post-nap than for items only remembered pre-nap (p < 0.01; Fig 3G).

### Post- and pre-cue spindle power positively and negatively predict subsequent memory

Recent evidence has suggested that post-TMR cue spindle power positively predicts memory, an effect that is maximal over the centroparietal midline electrode, CPz (Antony et al., 2018a, 2018b). We therefore asked whether these signals predict subsequent memory for items that were remembered pre-nap. We calculated spindle power by bandpass-filtering the raw EEG data between 11-16 Hz and then finding the root-mean-squared (RMS) value at each timepoint using a moving window of ± 100 ms. We found that post-cue spindle power was significantly higher from 1000 to 4500 ms for later-remembered than later-forgotten targets (later-remembered: 0.016 ± 0.06; later-forgotten: −0.74 ± 0.36, *t*(22) = 2.15, *d_z_* = 0.45, *p* = 0.043). Note that this analysis did not reach significance using only the early interval (1000-1500 ms) that we chose *a priori* based on previous studies, as described in the *Methods* (later-remembered: 0.69 ± 0.17; later-forgotten: −0.05 ± 0.48, *t*(22) = 1.49, *d_z_* = 0.31, *p* = 0.15).

Intriguingly, while post-cue spindle power may indicate that successful reactivation has occurred, pre-cue spindles may prevent reactivation because cues fall within the spindle refractory period (Antony et al., 2018b). Therefore, we next asked whether pre-cue spindle power over CPz negatively predicted subsequent memory for items that were remembered pre-nap. Note that, for this analysis, we did not perform standard baseline correction, as spindle power within the baseline interval is our variable of interest. Indeed, pre-cue spindle power was significantly lower from −2000 to 0 ms for later-remembered than later-forgotten items (later-remembered: 3.24 ± 0.16; later-forgotten: 3.73 ± 0.29, *t*(22) = 2.42, *d_z_* = 0.50, *p* = 0.024). In sum, both post- and pre-cue effects replicate previous findings and allow us to use spindle power over CPz as an independent predictor of lateralized reactivation evidence.

### Post- and pre-cue spindle power positively and negatively predict lateralized memory reactivation

We next asked whether post-cue spindle power was associated with improved lateralized memory reactivation and pre-cue spindle power associated with impaired lateralized memory reactivation. First, we took all trials where pre-nap memory was correct and split these trials by median post-cue sigma RMS values. We found significant lateralized memory reactivation for trials with above-median (*p* < 0.01), but not below-median (*p* = 0.3), RMS power (Fig 4A,B). Moreover, when we subtracted the AUC values between these conditions on each resampled subset (as above), the strength of lateralized memory reactivation differed significantly (*p* < 0.01; Fig 4C). Next, we took all trials where pre-nap memory was correct and split these trials by median pre-cue RMS values. As predicted, we found significant lateralized memory reactivation for trials with below-median (*p* = 0.01), but not above-median (*p* = 0.42), RMS power (Fig 4D,E). When contrasting high and low pre-cue spindle power directly, there was marginally higher lateralized memory reactivation in the low pre-cue RMS condition (*p* = 0.095; Fig 4F).

**Figure 4.**
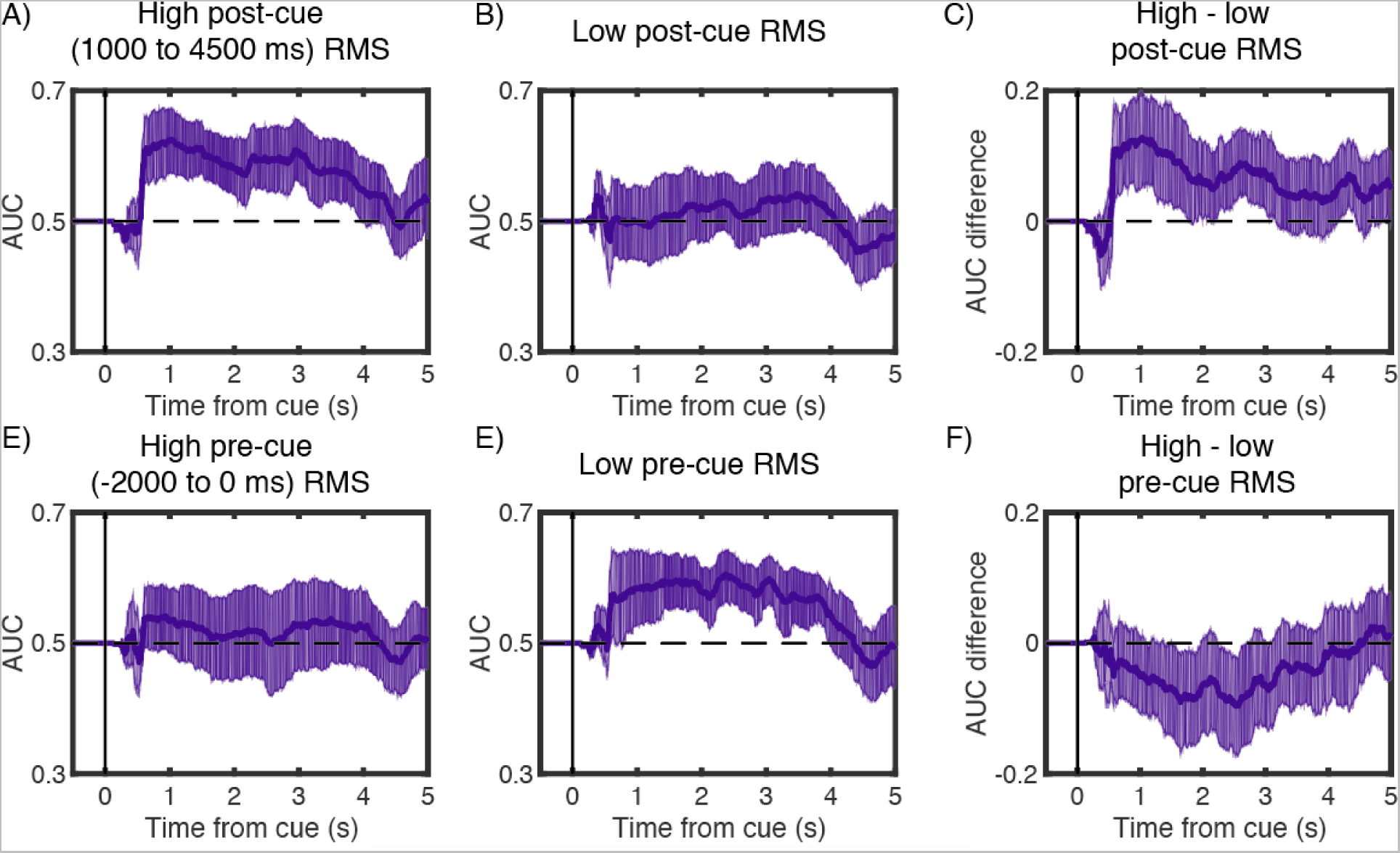
Content-related reactivation increased with higher post-cue spindle power and decreased with higher pre-cue spindle power. We plotted AUC-based classification performance as a function of post-cue (1000 to 4500 ms) and pre-cue spindle power (−2000 to 0 ms). Trials were split by the median RMS power in the sigma band of each participant. Classifier accuracy was significantly above chance on trials with above-median post-cue spindle power (A), but not below-median spindle power (B). Moreover, there were significant differences in classifier accuracy between these conditions (C). Conversely, classifier accuracy was not significantly above chance on trials with above-median pre-cue spindle power (D), but it was for trials with below-median spindle power (E), and there were marginally significant differences in classifier accuracy between these conditions (F). In all graphs, classification was performed using 1000-ms windows centered on each timepoint and error bars indicate the edges of the middle 95^th^ percentile of classification accuracies.

### Lateralized spindle incidence and power analyses

One report found that, after training participants to learn visual stimuli in a spatial array in either left or right hemisphere-heavy blocks, TMR cues elicited differences in the number and amplitude of lateralized spindles (Cox et al., 2014). Note that their analysis focused specifically on fast (> 13.5 Hz) spindles. Therefore, we tested whether these differences existed using a) both all lateralized channels pooled together and b) specifically using significant channels from the above study. For these analyses, we considered spindles that started between 0-4 s after the cue and were greater than 13.5 Hz in frequency. For spindle incidence, we performed double subtractions on the average number of spindles occurring over each electrode based on cue type and channel hemisphere [e.g., (number of spindles after left cues in C3 – C4) - (number of spindles after right cues in C3 – C4)]. For spindle amplitude, we averaged the power of each spindle (calculated as the sum of the absolute value of its data points) for all spindles within a particular condition [e.g., (mean spindle power after left cues in C3 – C4) - (mean spindle power after right cues in C3 – C4)]. Pooling all 27 lateralized channels together, we found no significant differences in the spindle incidence [mean subtracted difference: 0.012 ± 0.013 spindles, *t*(22) = 0.87, *d_z_* = 0.18, *p* = 0.40] or amplitude [15 ± 43 µV, *t*(22) = 0.34, *d_z_* = 0.07, *p* = 0.74]. For the specific channel analyses, we tried two channel pairs (TP7-TP8 and P7-P8). Neither produced significant differences in incidence [TP7-TP8 difference: 0.008 ± 0.04 spindles, *t*(22) = 0.22, *d_z_* = 0.05, *p* = 0.83; P7-P8 diff: 0.05 ± 0.03 spindles, *t*(22) = 1.43, *d_z_* = 0.30, *p* = 0.17] or amplitude [TP7-TP8 difference: 169 ± 85 µV, *t*(22) = 1.38, *d_z_* = 0.42, *p* = 0.20; P7-P8 diff: 9 ± 53 µV, *t*(22) = 0.13, *d_z_* = 0.04, *p* = 0.90]. Finally, these analyses were similarly not significant when considering all slow and fast spindles together (all *p* > 0.29).

## Discussion

We demonstrated that learning-related signals from individual episodic memories re-emerge when cued during sleep by decoding lateralized activity associated with specific sound cues. Evidence for reactivation was only significant if pre-nap memory was correct, suggesting this effect only occurs for learned information. Furthermore, reactivation evidence also predicted post-nap memory, specifically linking reactivation with memory stabilization. We also found post- and pre-cue spindle power positively and negatively predicted subsequent memory, respectively. Moreover, reactivation evidence was stronger for trials with high post-cue spindle power and low pre-cue spindle power (the former contrast was significant; the latter contrast was trending). Collectively, these analyses support the idea that sleep benefits memory retention by reinstating learning-related neural activity from individual learning episodes.

In our study, content-related differences in neural activity were detectable for more than four seconds after the cue. This result fits with results from other studies showing bouts of replay that extend for multiple seconds. For example, two studies using TMR in rodents found differences in learning-related neural activity for a number of seconds, lasting until the next cue arrived (Bendor and Wilson, 2012; Rothschild et al., 2017). While some human TMR studies have only reported neural differences predicting subsequent memory within the first ∼ 1.5 s (Schreiner et al., 2015; Lehmann et al., 2016; Farthouat et al., 2017; Antony et al., 2018b, 2018a; Belal et al., 2018), others have found differences up to and later than 2 s post-cue (Schreiner et al., 2015; Cairney et al., 2018). Importantly, this long window of reactivation may reflect brief bouts of reactivation occurring at variable times within the window, rather than continuous replay within the window.

To the best of our knowledge, our results are the first to show that, across trials, reactivation evidence on any particular trial increases with higher post-cue spindle power. These results nicely dovetail with findings showing that reactivation of neural content temporally coincides with spindles (Cairney et al., 2018). We also found a trend towards reactivation evidence decreasing with higher pre-cue spindle power; this converges with results showing that TMR cues are effective when followed by spindles and less effective when presented in the spindle refractory period, when ensuing spindles are unlikely (Antony et al., 2018b). Our results strongly support a recent mechanistic framework of memory reactivation proposing that memories are reinstated during spindle events and spindle refractoriness prevents further reinstatement of other memories (Antony et al., 2018c). However, we did not find evidence for the related idea that localized spindles support reactivation (Bergmann et al., 2012; Johnson et al., 2012; Cox et al., 2014). We hope future studies, perhaps using techniques with finer spatial resolution such as electrocorticography or magnetoencephalography or tasks engaging different areas of the neocortex, will clarify this issue.

One limitation of our study is that, although our cueing manipulation resulted in numerical differences between cued and uncued targets in the expected direction, this difference was not significant (when we limited the analysis to items that were remembered correctly pre-nap, the difference was marginal, *p* = .07). One possible explanation for this non-significant effect is that the pre-nap and post-nap tests differed in a way that was likely to engage different sorts of retrieval processing. The retrieval cue in each trial of the pre-nap test was the sound associated with the object (picture not shown), whereas the retrieval cue in each trial of the post-nap test was the item picture (the sound was not played). We omitted pictures in the pre-nap test to create fewer distractions for participants to perform motor imagery, and we omitted sounds in the post-nap test to ensure participants correctly associated visual items with locations and not sounds with locations, as we have done in previous studies (Antony et al., 2018a, 2018b). These differences in retrieval demands may have added noise to prepost comparisons, obscuring TMR effects.

In sum, our findings support the idea that sleep benefits memory by reactivating learning-related neural activity. Furthermore, reactivation following an auditory cue was linked with spindle occurrence, in keeping with our prior description of spindle refractoriness (Antony et al., 2018c). Memory reactivation was facilitated when the cue was followed by a spindle, as evidenced both by later memory performance and by EEG analyses. Moreover, this paradigm – using lateralized differences as a means of tracking reactivation of specific episodic associations – could be useful for uncovering nuanced relationships between reactivation and memory (Lewis-Peacock and Norman, 2014) and investigating how reactivation differs across physiological states (Andrillon et al., 2016; Jiang et al., 2017; Schönauer et al., 2017; Belal et al., 2018) and human populations (Mander et al., 2013; Westerberg et al., 2015).

## Methods

### Participants

Twenty-four participants (14 female, 18-33 years old, mean 22.3) with normal or corrected-to-normal vision and fluent in English were recruited via campus flyers and online scheduling software. Twenty-nine other participants were excluded for not sleeping long enough to receive at least one round of sleep cues. Participants were given hourly monetary compensation for participating ($20/hr) and small additional increases based on good performance (up to $5). To increase the likelihood of a successful nap, we requested that participants go to bed at their normal sleeping time but wake up two hours earlier than normal. We also requested that they abstain from alcohol the night before and coffee the morning of the experiment. Written informed consent was obtained in a manner approved by the Princeton University Institutional Review Boards.

### Experimental setup

Visual stimuli were presented on a 120 Hz LCD monitor in the testing room outfitted with an infrared touch screen frame. Touches on this display were registered as mouse input. Audio cues (during both wake and sleep) were played through a speaker system in the testing room.

### Stimuli

Participants viewed 128 images of common objects. Each image was shown in one of eight stimulus locations with a 300 x 300 pixel resolution. Sixty-four of these were associated with inherently related sounds (e.g. cow-[moo]) lasting up to 500 ms; the sounds were adapted from a previous study (Oudiette et al., 2013) and an online repository at www.freesound.org. Shorter sounds were looped, while longer sounds were trimmed, sped up, and pitch-corrected to keep them recognizable. During the nap, sleep cues were embedded in constant white noise (∼ 44 dB), resulting in increases of no larger than 5 dB.

### Design and procedure

The experiment included four phases: learning, pre-nap test, nap, and post-nap test (Fig 1). Participants arrived in the lab at 11AM and EEG electrodes were attached. They completed the learning phase task, took a 15-min break, and then took the pre-nap test. Following the test, participants napped in the testing room for 90 min from approximately 13:00 to 14:30. After the nap, participants were given a 90 min break during which they were allowed to leave the lab. Participants returned at 16:00 to complete the post-nap test.

In the learning phase, participants were trained to execute a lateralized motor action corresponding to the locations of 64 image-pairs. Each pair consisted of one target and one non-target image; the pair was presented along with an audio cue that was inherently associated with the target (e.g., cow-“moo”) – thus, the content of the audio cue (“moo”) specified to the participant which image was the target and (by exclusion) which image was the non-target. Non-targets had no immediately obvious sounds (e.g. avocado) and were included along with the target to ensure participants paid attention to the semantic content of the images.

Target - non-target pairs and image locations were assigned pseudorandomly for each participant. Images appeared in one of eight possible locations, each visually designed on the screen by a black, square outline. The target and non-target for a given sound were always on opposite sides of the screen. We assigned image locations such that each location was associated with 16 images, an equal number of which were targets and non-targets.

Participants underwent two blocks of image-location learning, each with 64 trials (one per image pair). Trial order was randomized in each block. A single trial comprised a fixation period, a stimulus presentation, an imagined selection, a physical selection, and a rest period. During the fixation period, participants were presented with a red central fixation cross and the eight empty image frames. After 1 s, an image-pair (target and non-target) appeared and the sound associated with the target was played. Participants had 5 s to imagine reaching out and touching the target with their corresponding hand (e.g., right hand for right side of screen). Then the fixation crosshair changed from red to green, signaling that the participant could execute their movement. If the participant touched the non-target or any other location, on-screen text instructed them to try again until they touched the target.

In the pre-nap test, each trial comprised a fixation period, a sound stimulus presentation, an imagined location selection, a physical location selection, and a rest period. After 1 s fixation, the target-related sound was played, and participants had 5 s in which to imagine touching the frame where the associated target had appeared during the learning phase, after which the crosshair turned green and they executed their movements. Each target was tested once, and participants received no feedback on response accuracy.

Following the test, participants napped for 90 min. All naps began between 12:00 and 13:00. Upon indications of SWS, we played half of the sounds associated with left targets (16) and half of the sounds associated with right targets (16) to reactivate their associated memories. Cues were separated by 4.5 s and cue order was randomized within loops. If the participant was asleep for 60 min and had not received cues, the experimenter loosened the criteria to include stage-2 sleep. The EEG was monitored continuously during cueing, and cues were stopped immediately upon indications of arousal; cues were resumed if the participant re-entered SWS. Participants were cued between 1-7 times per cue (mean = 3.32 ± 0.42) depending on how long they spent in SWS and stage-2 sleep. If by 90 min the participant was still asleep and had not completed a loop of cues, they were kept for an additional 20 minutes to try to reach this criterion. Table 1 shows a full breakdown of the stages and where cues fell with respect to those stages. Note that sleep stages are assigned based on whichever stage is more prevalent within each 30 s epoch, meaning that sounds can occur during unintended stages.

**Table 1.**
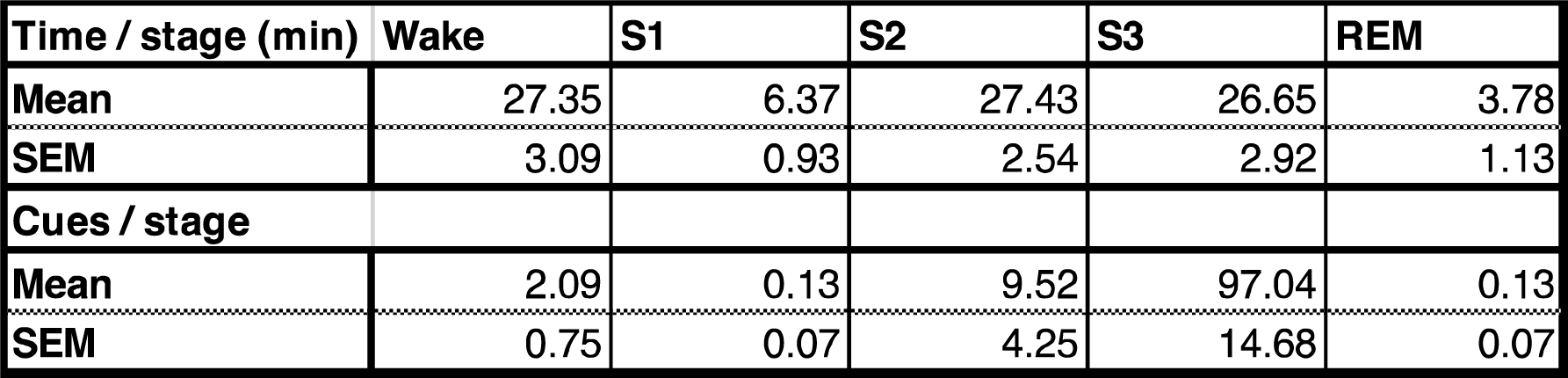
Time in each sleep stage and number of sounds per stage (min ± SEM).

After the nap, participants left the lab and returned again 90 minutes later for the final phase. They then completed a post-nap item-location test on both target and nontarget locations (128 trials total). On each trial, an image appeared in the center of the screen surrounded by the 8 empty frames as before. The participant was instructed to touch the frame where they believed the image appeared during the learning phase. After this test, participants were compensated for their time and left the laboratory.

### EEG data acquisition and preprocessing

Continuous EEG data were recorded during the learning phase, pre-nap test, and nap using Ag/AgCl active electrodes (Biosemi ActiveTwo, Amsterdam, The Netherlands). Recordings were made at 256 Hz from 64 scalp EEG electrodes from locations on the standard 10/20 layout plus intermediate 10% electrodes. Additionally, we placed two mastoid channels behind the left and right ears for offline re-referencing, a horizontal electrooculogram (EOG) electrode was placed next to the right eye, a vertical EOG electrode was placed under the left eye, and an electromyogram (EMG) electrode on the chin.

EEG data were processed using a combination of internal functions in EEGLAB (Delorme and Makeig, 2004) and custom-written scripts. Data were re-referenced offline to the average signal of the left and right mastoids, then they were high-pass filtered at 0.1 Hz and low-pass filtered at 60 Hz in successive steps. For each wake recording, we removed artifactual segments in the continuous EEG of the learning phase. We ran spatial independent components analysis (ICA; Delorme & Makeig, 2004) for each participant to remove eyeblink components. Any electrodes with artifacts for long stretches of time were marked and interpolated within-subject after ICA.

Next, we created across-subject components. The learning phase data were segmented into epochs including the 5-s imagination period with 2 s of baseline (i.e., −2 to 5 s relative to the imagination cue). We subtracted lateralized channels (e.g., C3-C4) and omitted channels along the midline to reduce the number of features from 64 channels to 27 channel pairs (Fig. 2). We then concatenated data from all 24 participants to create one across-subject dataset from which we ran across-subject spatial ICA. This procedure allowed us to reduce the dimensionality of the data while maintaining consistent features across individuals.

To assess spindle power, we first bandpass-filtered the raw signal between 11-16 Hz. Then we calculated the root-mean-square (RMS) value for every time point using a moving window of ± 100 ms (Antony et al., 2018b, 2018a). To assess whether there were significant differences in spindle power between memory conditions, we averaged RMS values across specified time windows for each participant and calculated differences across conditions using within-subject *t*-tests. Our specified pre-cue window was −2000 to 0, based on a previous study (Antony et al., 2018b). Our *a priori* post-cue window was 1000 to 1500 ms (Antony et al., 2018a), though we eventually widened this window to 1000-4500 ms after observing that left/right decoding accuracy stayed above chance up to 4.5 s after cue onset.

### Sleep physiological analyses

Sleep stages were determined by an expert scorer according to standard American Academy of Sleep Medicine (AASM) criteria (Rechtschaffen and Kales, 1968). Artifacts (large movements, blinks, arousals, and rare, large deflections in single channels) during sleep were identified visually and rejected in five-second chunks following sleep staging.

### Classification during wake for feature selection

We first asked how well each component discriminated between left and right imagination conditions during wake in two successive steps. First, we performed baseline correction on each trial by subtracting out the average activity from −1000 to 0 ms relative to the onset of the visual stimulus. We divided each trial’s data into overlapping time bins of length 1000 ms; for each time bin, we trained a single-feature, logistic regression classifier (that component’s average value across the time bin) to discriminate between left and right trials, using a leave-one-subject-out cross-validation procedure. This procedure yielded an area under the curve (AUC) value for each participant and each time bin. We then used a bootstrap procedure to generate 95% confidence intervals for the average AUC value (across participants) for each time bin. For this procedure, we resampled participants’ AUC scores with replacement 1000 times. For each resampling, we computed the average AUC across (resampled) participants, yielding a bootstrap distribution of 1000 average-AUC scores for each time bin. After sorting the AUC values at each time bin from low to high, we counted the number of time bins for which the middle 95th percentile of bootstrapped trials passed above or below chance (at the 2.5th or 97.5th percentile); we refer to this measure as “number of significant time bins.”

Second, to correct for multiple comparisons (i.e., looking at multiple time bins), we scrambled the left and right condition labels 200 times and repeated the bootstrap procedure above for each of the scrambles. This allowed us to compute a null distribution on the “number of significant time bins” measure described above. If, for a given component, the number of individually significant time bins exceeded 95% of the null distribution, the component was deemed to significantly discriminate between left and right trials (corrected for multiple comparisons).

### Classification during sleep

To compute classification accuracy during sleep (in response to TMR cues), we computed the time series of activity (on each trial, in response to the TMR cue) of each of the five components that were identified as significant using the wake-classification procedure outlined above. We again performed baseline correction from −1000 to 0 ms relative to cue onset. As with the wake-classification procedure, we divided each trial’s data (time locked to the TMR cue) into overlapping time bins of length 1000 ms, and we trained a separate classifier for each time bin. Putting all of this together, each of these time-bin-specific classifiers was fed a 5-dimensional vector for each TMR cue (reflecting the average activity of each of the five components during the relevant time bin, in response to that cue); the job of the classifier was to determine whether the TMR cue that was just played was associated with a left- or right-sided movement. As with wake classification, we used a leave-one-subject-out cross-validation procedure that yielded an AUC score for each time bin for each participant. For the sleep data, we used an L1-regularized logistic regression classifier (λ = 0.1). Multiple-comparisons-corrected significance was assessed using the same bootstrapping-and-scrambling procedure that we used for the wake classifier. Note that, in some cases, the regularization was strong enough such that all of the feature weights were zero, resulting in a classification score that was exactly at chance. Note also that, for the sleep-EEG analyses, we omitted one participant who did not have any items that were initially remembered and later forgotten (making it impossible to include them in subsequent-memory analyses). We did this to ensure that all of our sleep-EEG analyses included the same participants, thereby maximizing comparability across these analyses. After this 24^th^ participant was dropped, we were left with 23 participants for all of the sleep-EEG analyses (importantly, the qualitative nature of our results was the same regardless of whether the 24^th^ participant was included; for example, re-doing the classifier analysis shown in Fig. 3A with all 24 participants still produced a significant result of *p* < 0.01).

First, we ran this sleep-classification analysis on all trials. Second, we filtered the data based on whether items were correct before the nap or not, hypothesizing that only items correct before the nap could be reactivated during sleep. Note that, for this and all other “filtered” analyses, we used the classifier that was trained on all trials, and we segregated test trials depending on whatever variable we were using to filter the data (here, pre-nap accuracy). We also ran an analysis to look at items that were remembered correctly pre-nap, and additionally filtered by whether items were remembered (or not) post-nap, hypothesizing that the content-related signal may predict subsequent memory. For the latter two ways of splitting the data, we also conducted subtraction analyses, whereby we calculated differences in the AUC (e.g., between pre-nap-correct and pre-nap-incorrect) after each iteration of resampling the data, and then we assessed whether these AUC differences were significantly different from zero using the same bootstrapping-and-scrambling procedure as above. The mean number of trials per subject that entered our analysis (omitting those falling during rejected artifacts) was as follows: items remembered on both the pre- and post-nap tests, 48.83 ± 15.4; items remembered on the pre-nap test, but not the post-nap test, 9.39 ± 4.14; items not remembered on the pre-nap test, but remembered on the post-nap test, 20.17 ± 7.34; items not remembered on either test, 27.61 ± 10.82. Note that this counts the same cue presented to the same subject (e.g., “moo” on the first and second round of cues) as separate trials.

We additionally investigated whether pre- and post-cue spindle power modulates this content-related signal. For these analyses, we calculated AUC as above after splitting trials by each participant’s median RMS power value in the pre-specified time windows (pre: −2000 to 0 ms; post: 1000 to 4500 ms). Similar to the above approaches, we also calculated subtracted differences between AUC values for trials above and below their corresponding median RMS values. Note that for these analyses, we only used trials that were correct before the nap.

## Data availability

The datasets generated during and/or analyzed during the current study and the analysis code will be made available at the time of publication on the Open Science Framework website.

Contributions
JWA, SL, KAP, and KAN conceived the experimental design. SL and PPB collected the data. BW, JWA, and KAN analyzed the data. BW performed the classification analyses. BW, JWA, and KAN wrote the manuscript. All authors discussed the results and revised the manuscript.

## Acknowledgements

Parts of this paper were adapted from SL’s undergraduate thesis. This work was supported by the CV Starr fellowship to JWA and the NSF BCS grant 1533511 to KAP and KAN.

